# Improving nanobody structure prediction with self-distillation

**DOI:** 10.64898/2025.12.01.691162

**Authors:** Montader Ali, Matthew Greenig, Mateusz Jaskolowski, Mia Crnogaj, Eva Smorodina, Haowen Zhao, Victor Greiff, Pietro Sormanni

**Affiliations:** Yusuf Hamied Department of Chemistry, University of Cambridge, Cambridge, UK; {,,,, |; Department of Immunology, University of Oslo and Oslo University Hospital, Oslo, Norway; {, |

## Abstract

Nanobodies are increasingly attractive therapeutic and biotechnological molecules, yet accurate structure prediction of their highly variable H-CDR3 loops remains a central challenge for machine learning models. Here, we investigate whether nanobody-specific structure prediction can be improved through curated synthetic data strategies. We systematically evaluate different data augmentation regimes, including self-distillation from unlabelled VHH sequences. To ensure structural plausibility of synthetic training samples, we develop **NanoKink**, the first sequence-based classifier of kinked versus extended H-CDR3 conformations, and apply stringent filtering criteria for non-canonical disulfide bond placement and conformational accuracy. On a curated benchmark enriched for challenging nanobody features, we show that, for a fixed training compute budget, a nanobody-specific model trained with filtered synthetic data significantly improves over baseline models and NanobodyBuilder2, achieving lower mean H-CDR3 RMSD and fewer structural violations, while remaining competitive with AlphaFold3 at approximately two orders of magnitude lower per-structure inference time. Our results highlight promising directions in synthetic data generation for nanobody structure modelling and provide a practical framework for optimisation of VHH structure prediction models.

## 1 Introduction

Nanobodies are single-domain antibodies naturally produced in camelids [1, 2]. These molecules are becoming increasingly recognised as a promising molecular modality for a wide range of therapeutic applications [3] following the approval of the first nanobody drug in 2019 [4]. While conventional antibody variable regions comprise two separate protein chains (VH and VL) that form a complex, nanobody variable regions contain only an isolated heavy chain (VHH). Through natural evolution of the adaptive immune system in camelids, this key difference has endowed nanobodies with unique structural properties, some of which also happen to improve their prospects as molecular tools in biotechnology for some applications. For instance, in conventional antibodies, the heavy/light chain complex is stabilised by hydrophobic interactions at the interface between the two chains [5]. However, in the absence of a light chain, evolution has produced hydrophilic substitutions at these interface positions in VHH domains [6], improving the expression yield, solubility, and thermostability profiles of nanobodies compared to typical paired antibodies [2, 7]. The VH/VL complex in conventional antibodies also generally produces a binding interface which is approximately flat or concave [2], restricting the epitope geometries that these molecules can target. Nanobodies, on the other hand, can also adopt a convex paratope, and hence are capable of targeting epitopes inaccessible to conventional antibodies [7]; in particular, the lack of a light chain can allow a nanobody’s complementarity determining region (CDR) loops to access concave epitopes like enzyme active sites [2, 8].

These properties have made nanobodies an attractive and popular target for antibody design campaigns, including in the emerging field of *in silico* design [9]. Recently, significant progress has been made towards the objective of *de novo* epitope-specific nanobody design using generative models, where the target protein and the desired binding site are provided as input to a model that conditionally samples nanobody sequences (and typically structures) that are predicted to bind the target at that site [10, 11, 12]. The broader community of ML-driven protein binder design has established that performing *in silico* screening of designed binders using orthogonal structure prediction models drastically improves experimental success rates [13, 14]. In this setting, only designs whose generated structures match an independent structure prediction are taken forward for experimental validation.

Thus, the ability to rapidly and accurately screen generated nanobodies to identify which are unlikely to fold into their designed monomeric structure would improve the prospects of generative design workflows. More broadly, access to accurate structure prediction tools greatly facilitates efforts to engineer the functional and biophysical properties of nanobodies. While AlphaFold3 is widely regarded as the state-of-the-art prediction model for bound and unbound nanobody structures [15, 16, 17], it is not available for commercial use and incurs significant runtime costs. Indeed, the runtime requirements of AlphaFold3 and its predecessor AlphaFold2 [18] have motivated the development of smaller antibody-specific models that operate at a fraction of the cost, many of which achieve comparable performance (on antibodies) to the AlphaFold series [19, 20, 21].

Yet for ML-driven nanobody structure prediction, significant challenges remain. The first is data availability: the PDB contains only around 2000 entries with nanobody structures, many of which are redundant. Nanobodies also have structural idiosyncracies - especially related to the H-CDR3 loop - that make them particularly challenging to model. The H-CDR3 loop in conventional antibodies is difficult to model accurately due to its sequence diversity and unique structural features [22], and nanobodies are no exception; in fact, nanobodies tend to have *longer* H-CDR3 loops than standard antibodies [2], making accurate structure prediction an even greater challenge. Since longer H-CDR3 loops may incur greater entropic costs on binding, many such nanobody loops also contain a unique structural feature known as a *non-canonical disulfide bond* (NCDB): a disulfide bond that forms between a cysteine residue on the H-CDR3 and another cysteine located elsewhere in the nanobody, acting to stabilise the loop and reduce its conformational space [23]. Since structure prediction models are trained with continuous-valued objectives, the “hard” geometric constraint of the NCDB is challenging to model accurately, despite its crucial function role. Furthermore, nanobody H-CDR3 loop structures tend to cluster into two conformational groups: *kinked* and *extended*, which are qualitatively distinct (see Figure 1) but challenging to classify exactly, even using experimental structures [24, 25]. Taken together, these features help to explain the observed performance gap in H-CDR3 structure prediction between conventional antibodies and nanobodies [19].

**Figure 1.**
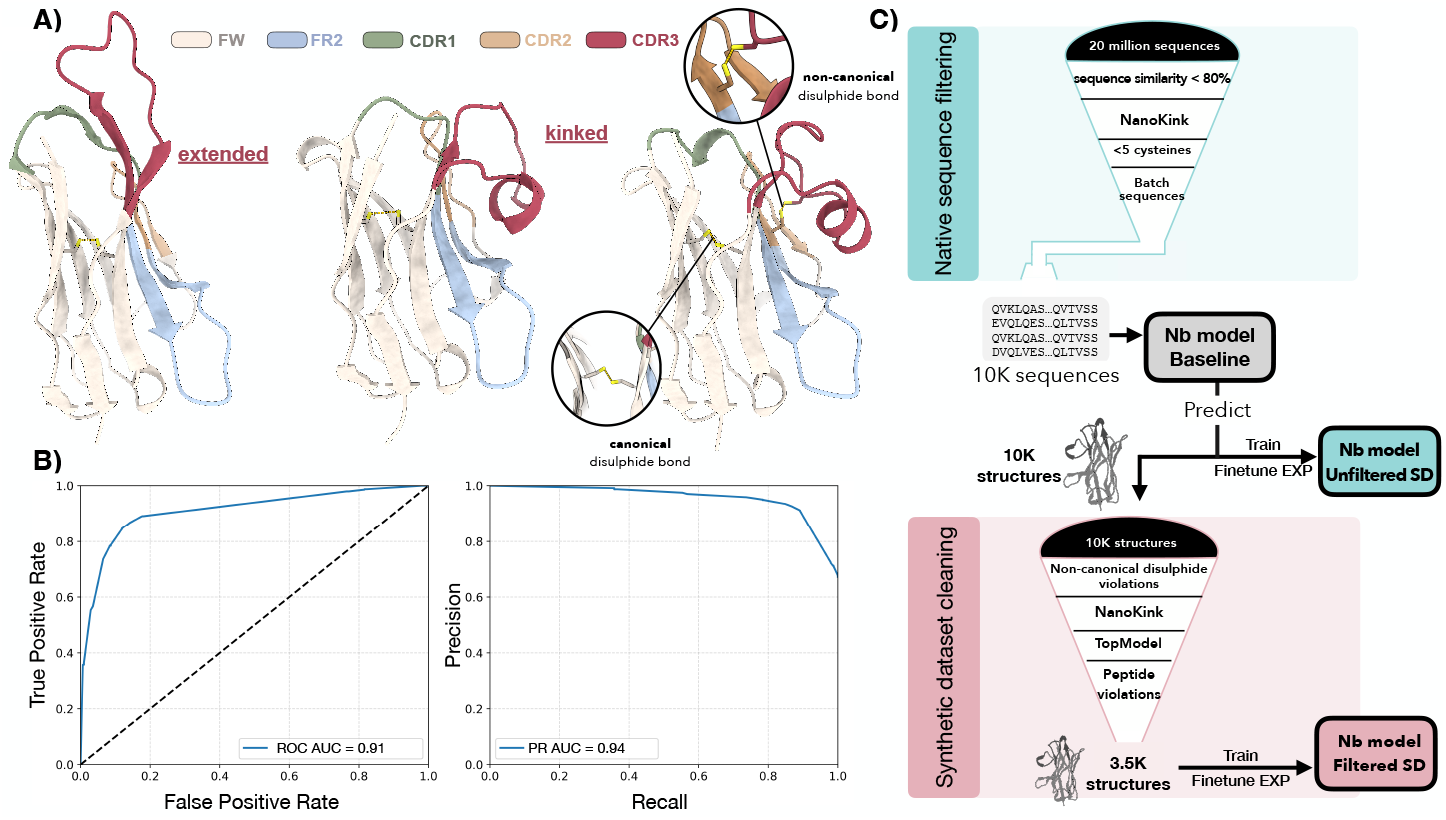
Nanobody-specific approaches to structure prediction. **A)** Unique features of nanobody H-CDR3 loops: extended (left) and kinked (middle) conformations, as well as non-canonical disulfide bonds (right). **B)** The NanoKink score achieves high prediction accuracy in classifying H-CDR3 loop conformations (kinked vs. extended) from sequence information alone. **C)** The Nanobody model’s self-distillation (SD) pipeline. First, a baseline model is trained on true VHH structures from SAbDab. Then, the baseline model is used to predict VHH structures from a large, unlabelled corpus of sequences. These sequences are then either used for training directly (“unfiltered”) or filtered based on structural violations, plausible NCDB geometry, and predicted H-CDR3 loop structures that match the conformational category predicted by NanoKink.

## 2 Our contribution

We sought to investigate if the performance of nanobody-specific structure prediction models could be improved with synthetic data curation strategies. One such strategy is *self-distillation*, a technique in which a trained model is used to make predictions on unlabelled data, which are then treated as ground-truth labels in another round of training for a newly initialised model [26, 27]. In AlphaFold2, this technique was demonstrated to robustly improve structure prediction performance (using predicted structures as training data) [18], and more recently it was applied in training the pan-immunoglobulin prediction model Ibex [21]. Nevertheless, best practices for self-distillation - and synthetic data sets more broadly - in the context of nanobody structure prediction remain largely unexplored. We thus set out to systematically evaluate a variety of design choices in the synthetic data regime for nanobody structural modelling, including the use of self-distilled structures and conventional antibody VH chains as synthetic training data.

To explore the effects of different configurations, we train a simple structure prediction model from existing architectural components. We benchmark our model against AlphaFold3 [15] and NanobodyBuilder2 [19] on a novel test set of 47 nanobodies that are selected to overrepresent structures with more challenging features such as longer H-CDR3 lengths and non-canonical disulfide bonds (Table 1). While we had also hoped to evaluate against the recently-published Ibex model [21], we could not obtain publicly available data on the exact structures included in their training set and opted not to evaluate on the NanobodyBuilder2 test set they used, due to its poor representation of the NCDB (Table 1). In addition to benchmarking structure prediction models using standard RMSD metrics, we also evaluate the *physical plausibility* of predicted nanobody structures based on structural violations, H-CDR3 loop conformational category, and NCDB prediction.

**Table 1.** Summary statistics of different VHH datasets, including the mean and standard deviation of H-CDR3 length, frequency of non-canonical disulfide bonds, and frequency of kinked H-CDR3 loop conformations. Our test set has no overlap with NanobodyBuilder2’s, and is chosen to overrepresent longer H-CDR3 loops and non-canonical disulfide bonds, allowing performance in these challenging settings to be accurately assessed.

| Dataset | H-CDR3 length (mean ± std) | NCDB (%) | Kinked (%) |
| --- | --- | --- | --- |
| SAbDab (all VHH chains) | 14.4 ± 4.7 | 17.0 | 69.6 |
| SAbDab (unique VHH sequences) | 14.3 ± 4.6 | 13.9 | 68.6 |
| NanobodyBuilder2 test set | 14.0 ± 4.5 | 2.5 | 80.0 |
| Nanobody model test set | 17.8 ± 4.3 | 31.3 | 62.5 |

Since state-of-the-art nanobody structure predictors still often fail to achieve atomistic accuracy [19, 21] - particularly in the H-CDR3 loop - we also hypothesised that it would be beneficial to use stringent filtering criteria when constructing self-distilled training sets to ensure that synthetic data samples were of sufficient quality to provide a signal that generalises to experimental nanobody structures. To that end, we introduce and evaluate a pipeline for filtering predicted structures to ensure they recapitulate the key features of natural nanobodies, such as accurately-placed non-canonical disulfide bonds and correct H-CDR3 conformations. We develop a nanobody H-CDR3 conformation classifier called **NanoKink** - the first method capable of classifying nanobody H-CDR3 conformations as *kinked* or *extended* from sequence information alone - and use it to screen self-distilled samples whose loop conformation is incorrectly predicted. Our results demonstrate that controlling the composition of synthetic training data notably improves structure prediction accuracy in terms of both RMSD and physical plausibility.

## 3 Methods

### 3.1 Model architecture

We adopt the architecture and training objectives of ABodyBuilder3 [20, 28] to train nanobody structure predictors (details in Appendix B). To incentivise correct placement of disulfide bonds, we implement an additional violation loss on the SG (sulfur) atom coordinates for cysteine residues. For a nanobody with N = 2, 3, or 4 cysteine residues, let ℬ ⊆{(*i, j*): 1 ≤ *i < j* ≤ *N*} denote the subset of cysteine residue index pairs involved in a disulfide bond (either canonical or non-canonical) and ℬ^*′*^ denote its complement, i.e. all pairs of cysteines that are non-bonded. The disulfide loss is then a hinge-style objective on the distances *d*_*ij*_ between predicted SG atom coordinates:

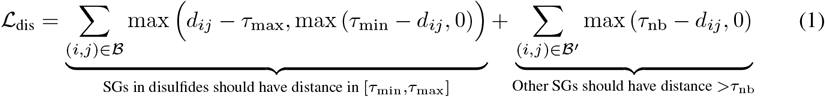

Where the thresholds were determined by examining bonded and unbonded cysteine distances in SAbDab [29]; we chose values of *τ*_min_ = 1.62, *τ*_max_ = 2.55, and *τ*_nb_ = 4.55.

### 3.2 Data sets

All VHH sequences in SAbDab [29] were clustered with MMSeqs2 at 100% sequence identity [30] and cluster assignments were used to obtain train/validation/test splits, avoiding data leakage from different PDBs with the same sequence. The test set was specifically curated to test the model’s capabilities over diverse VHH structural features, including a balanced panel of *kinked* and *extended* H-CDR3 conformations, 14 structures with non-canonical disulfide bridges, and a distribution of H-CDR3 lengths that is deliberately longer than that observed in natural VHH PDBs, to stress-test robustness on long loops (Table 1). A baseline model was trained on these experimental VHHs. We also used these structures to define the NanoKink scoring function, which uses position-specific amino acid frequencies to estimate the probability of an H-CDR3 loop being kinked or extended from sequence information alone (details in Appendix A).

A VHH sequence database was derived from the camelid repertoire compiled by Ramon et al. [31]. After applying our quality-control and filtering steps, this yielded a set of 20 million VHH sequences, which was clustered using MMseqs2 at a threshold of 80% sequence similarity, removing singletons.

The remaining sequences were then evaluated with NanoKink. Only sequences with NanoKink scores below 0.2 or above 0.99 were retained, ensuring inclusion of sequences predicted with high confidence to adopt either an extended or kinked conformation. Sequences exhibiting atypical cysteine counts (fewer than 2 or greater than 5) were excluded. The resulting dataset of 160,000 sequences was partitioned into 16 subsets of 10,000 sequences each, stratified to preserve the original distributions of CDR3 length, NanoKink score, and cysteine counts. A single random subset was selected for the synthetic data experiments presented here. Using our baseline model trained on SAbDab VHH structures, we predicted the structures for these 10K VHH sequences and filtered them according to NanoKink score, non-canonical disulfide bond placement, and structural violations (details in Appendix C). The synthetic data sets of 10K unfiltered structures and 3.5K filtered structures were used for pre-training separate Nanobody models before finetuning on experimental VHH structures.

## 4 Results

We trained different versions of a Nanobody model using a variety of synthetic and experimental structure sets and evaluated the trained models using our test set (Table 2), benchmarking our approach against AlphaFold3 (AF3) [15] and NanobodyBuilder2 [19] on the same stratified test set. For AF3, we ran prediction using both multiple sequence alignments and structural templates, as well as without. Strikingly, we observed an immense reduction in performance in the MSA- and template-free regime; upon visual inspection, many of these high-RMSD predictions suffer from significant errors in the framework prediction (higher even than CDRs 1 and 2, see Table 3). This observation suggests that evolutionary and/or template information for AF3 is indispensible for accurately modelling VHH frameworks, ultimately leading to more accurate CDR predictions.

**Table 2.** Structure prediction performance metrics for different configurations of the Nanobody model and other nanobody structure prediction models. From left to right: **mean H3 RMSD** (C*α* RMSD of the H-CDR3 loop after superimposing all VHH C*α* atoms- see Figure 2 for distribution), **NCDB** (% of true non-canonical disulfide bonds correctly predicted), **K/E** (% of predicted H-CDR3 loop structures that adopt the same conformational category as the true structure, kinked/extended), and **Violations** (structures with any residues with non-planar amide bond geometry, any adjacent C*α*-C*α* distance *>*4Å, and/or any peptide bond length *>*2.5Å). For AlphaFold3, scores are shown for predictions made using a multiple sequence alignment and structural templates, and those made without either (in brackets). The highest-scoring value for each metric is shown in **bold**, with the second-highest scoring value underlined.

| Model | H3 RMSD (Å) ↓ | NCDB (%) ↑ | K/E (%) ↑ | Violations (%) ↓ |
| --- | --- | --- | --- | --- |
| Nb model (VHH) | 3.57 | 21.4 | 93.6 | 74.5 |
| Nb model (VH + VHH) | 3.44 | <u>57.1</u> | 93.6 | 74.5 |
| Nb model (filtered SD) | <u>3.28</u> | 50.0 | <u>95.7</u> | <u>42.6</u> |
| Nb model (unfiltered SD) | 3.42 | 28.6 | **97.9** | 76.6 |
| AlphaFold3 | **2.82** (7.98) | **92.9** (35.7) | **97.9** (76.6) | **2.1** (25.5) |
| NanobodyBuilder2 | 3.58 | 28.6 | 93.6 | 80.9 |

**Table 3.** Detailed C*α* RMSD statistics for the Nanobody (Nb) model trained on different datasets, and other structure prediction models. We report the RMSD for CDR1, CDR2, CDR3, all framework regions (FWs), and the entire nanobody (all) after superimposing structures using the C*α* atom positions of the entire VHH chain. Regions are determined based on the AHo numbering scheme. AlphaFold3 values in brackets represent runs without MSA and templates.

| Model | CDR1 (Å) ↓ | CDR2 (Å) ↓ | CDR3 (Å) ↓ | FWs (Å) ↓ | All (Å) ↓ |
| --- | --- | --- | --- | --- | --- |
| Nb model (VHH) | 1.83 | 1.42 | 3.57 | 1.00 | 1.70 |
| Nb model (VH + VHH) | 1.79 | 1.24 | 3.44 | 0.91 | 1.64 |
| Nb model (filtered SD) | 1.74 | 1.29 | 3.28 | 0.94 | 1.59 |
| Nb model (unfiltered SD) | 1.81 | 1.40 | 3.42 | 0.99 | 1.67 |
| AlphaFold3 | **1.51** (5.54) | **1.10** (5.07) | **2.82** (7.98) | **0.78** (6.31) | **1.39** (6.73) |
| NanobodyBuilder2 | 1.86 | 1.37 | 3.58 | 0.92 | 1.69 |

Table 2 also shows that augmenting the Nanobody model with additional data - both antibody VH chains and synthetic VHH structures - improved the performance of the baseline model, both in terms of RMSD of the H-CDR3 loop and correct NCDB placement. Within this fixed compute budget, the most consistently performant regime used pre-training on a filtered synthetic dataset and finetuning on experimental VHH structures (“filtered SD”), which produced statistically significant improvements over both the baseline Nanobody model and NanobodyBuilder2 (Figure 2 in Appendix D) in terms of H-CDR3 RMSD (*p <* 0.05, Wilcoxon’s signed rank test) and strong results across all physical plausibility metrics. Notably, filtered self-distillation was the only data augmentation regime that managed to substantially reduce the incidence of structural violations.

**Figure 2.**
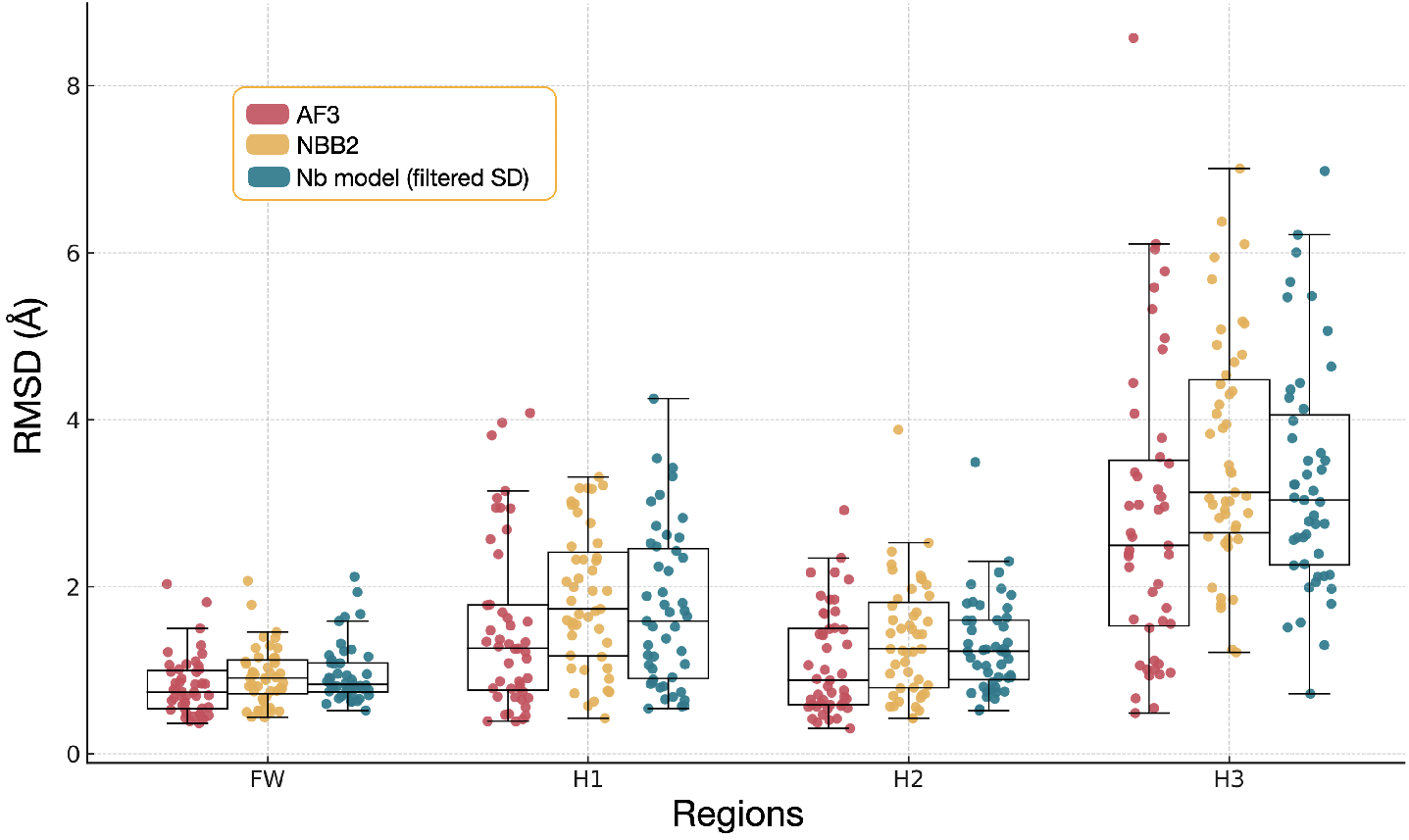
Swarmplot of the RMSD (Å) of C*α* coordinates for each nanobody region, after superimposition of the C*α* coordinates. The best H3 model from the Nanobody models is shown (filtered SD), alongside the NBB2 and AF3 models.

## 5 Conclusion

Accurate prediction of the nanobody H-CDR3 loop from sequence information alone is a powerful tool for nanobody engineering and design, but achieving atomistic accuracy remains a challenge. As a large foundation model for biomolecular structure prediction, AlphaFold3 achieves state-of-the-art performance across the board on this task (Table 2, Figure 2) [15]. However, AF3 is bottlenecked by its reliance on MSAs and structural templates, both of which increase runtime but are indispensable for prediction accuracy (Table 2 and Table 4). In the context of large-scale screening of sequence libraries - for instance, those generated in *de novo* design campaigns [10, 11, 12] - model runtime is a key consideration when screening millions of structures, and this Nanobody model is superior to AF3 in terms of runtime on both CPU and GPU (Table 4). While we do not explicitly compare to Ibex [21] due to non-overlapping test sets, we note that they reported a similar mean H3 RMSD improvement over NanobodyBuilder2 of 0.19. Within this fixed compute budget setting, we report similar performance gains simply by modulating the composition of synthetic data used for pre-training, without any architectural improvements. Combining our approach with better optimised model architectures [21], as well as training until convergence, may yield even more effective nanobody structure prediction models.

**Table 4.** Wall-clock prediction time for AlphaFold3 and the Nanobody model on the test set of 47 nanobodies used in this study. For AlphaFold3, values in parentheses correspond to predictions that exclude multiple sequence alignment (MSA) and structural template searches. All values are averaged over three independent runs. All timings were measured with a NVIDIA Quadro RTX 8000 GPU and a AMD EPYC 7702 CPU.

| Model | Total time (s; 3 runs) ↓ | Per-structure time (s) ↓ |
| --- | --- | --- |
| AlphaFold3 (GPU) | 2676.9 (1841.8) | 57.0 (39.2) |
| Nanobody model (GPU) | **19** | **0.4** |
| Nanobody model (CPU) | 39.4 | 0.8 |

## Appendix A NanoKink

After filtering low-resolution (*>*4Å) and outlier structures from SAbDab (using framework C*α* RMSD *>*5Å relative to a reference), each nanobody structure was assigned a binary label denoting whether its H-CDR3 loop adopts a kinked conformation. We initially evaluated a geometric angle-based approach [24] but observed frequent misclassifications upon visual inspection. Because kinked CDR3 loops tend to contact framework 2 (FR2), we defined a structure as “Kinked” when any residue within the CDR3 loop (AHo definition; excluding two stem residues on each side) formed a heavy atom contact (*<*4.5Å) with any FR2 residue; otherwise it was labelled “Not Kinked”. Visually, this contact-based criterion produced more reliable labels than the angle-based method.

Using the structurally-labelled sets, we generated AHo-numbered multiple sequence alignments separately for kinked and non-kinked sequences and tested for amino-acid enrichment at each position. For every AHo position and amino acid, we built a 2×2 contingency table of sequence counts and applied a two-sided Fisher’s exact test to assess association between the presence of the amino acid and membership in the kinked/non-kinked MSAs (excluding gaps). P-values were adjusted for multiple testing using the Benjamini–Hochberg procedure, and significance was assessed at FDR-adjusted p < 0.05 (q < 0.05). Positions for which any amino acid met this threshold were designated as “hallmarks”. Log fold change (LFC) scores were calculated at these hallmark positions for a large nanobody sequence database (∼ 22 million unique VHH sequences) derived from the camelid VHH repertoire assembled by Ramon et al. [31]. Prior to scoring, we filtered the database to sequences containing exactly two cysteines at the canonical positions, yielding 10,157,265 sequences. For a given amino acid *a*, the LFC is 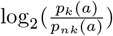, where *p*_*k*_(*a*) is the empirical frequency of *a* in the kinked MSA and *p*_*nk*_(*a*) is its frequency in the non-kinked MSA. We computed a score for each sequence as the sum of these LFCs over all hallmark positions and labeled sequences as kinked (score > 2.66) or non-kinked (score < 0). A second round of this analysis was then performed on the broadened sequence database, yielding new MSAs for the kinked/non-kinked groups and a new set of hallmark positions. LFC scores at these hallmarks (computed on the expanded database MSAs) are used by the final predictor.

Rather than summing over raw LFC values, we found it was beneficial to shrink the LFCs via a saturating transform before summation:

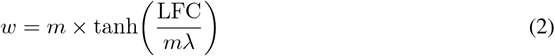

where *m* is a scaling factor controlling the maximum transformed LFC (default *m* = 10.0) and *λ* is a shape factor (default *λ* = 1.0). This transformation bounds extreme LFC values while retaining sign and relative magnitude. The final raw score for an input VHH sequence is the sum of weights over all hallmark positions.

Raw scores were mapped to calibrated probabilities in [0, 1] using an isotonic regression model trained on a subset of the structural dataset. We evaluated isotonic regression against Platt scaling via 5-fold cross-validation and selected isotonic regression based on a lower Brier score (0.1099 vs 0.1227). The deployed calibrator was trained on 750 unique nanobody structures restricted to sequences bearing only canonical cysteines.

NanoKink will be available soon via our web server, with the code provided on GitHub.

## APPENDIX B Nanobody model details

Like ABodyBuilder3 [20], the Nanobody model uses a single network with eight AF2 structure module layers [18, 28] (randomly initialized), with input single features as a one-hot encoding of the amino acid sequence and input pair features as a one-hot encoding of the sequence positional offset between residues (clamped at a maximum of 32). Single features are updated with interleaved invariant point attention and transition blocks, while pair features are updated only with transition blocks. An embedding dimension of 128 is used for both the single and pair features, producing a model with 8.3M parameters. Training hyperparameters are taken from ABodyBuilder3; we use the RAdam optimizer [32] with a batch size of 32 and a cosine-annealed learning rate schedule with warm restarts [33]. All-atom FAPE loss is computed at every layer of the structure module, as well as AF2’s pLDDT and structural violation losses on the coordinates output by the final layer.

We performed all training and finetuning runs for 24 hours on a single H100 GPU, corresponding to approximately 100K optimisation steps for each configuration. This protocol equalises the number of optimisation steps and wall-clock compute across synthetic data regimes, so all comparisons should be interpreted as evaluating performance under a fixed compute budget. Some configurations may not have fully converged within this budget, and training the Nanobody models to complete convergence may result in better performance, an exploration we leave to future work.

## APPENDIX c Synthetic sequence filtering

We performed stringent filtering on the structural properties of the 10K structures predicted by our baseline model to obtain our filtered synthetic data set. First, we filtered for structures in which all disulfide bonds - both canonical and non-canonical - were correctly placed. Pairwise distances between SG atoms were computed to identify canonical disulfide bonds between residues at positions 23 and 106 (AHo numbering) as well as potential non-canonical disulfide bonds, using a 2.5Å distance cutoff. Additional rules were applied to flag odd cysteine counts and violations of an isolation criterion, whereby unpaired cysteines located within 5Å of paired residues were considered atypical. Furthermore, incorrectly predicted H-CDR3 conformations that did not align with NanoKink’s score were removed. For this, we defined framework region 2 (FR2) using AHo positions 43–56 (inclusive on both sides), and the CDR3 loop segment as AHo 108–138 with the stem residues 108, 109, 137, and 138 excluded (to avoid confounding kinkedness with stem packing structures). For each structure, all residue–residue pairs between this CDR3 segment and FR2 were evaluated for heavy-atom proximity, and structures were classified as kinked if at least one unique CDR3–FR2 heavy-atom contact (*<*4.5Å) was present; structures with zero such contacts were classified as extended. Finally, structures that produced TopModel’s non-planar bond violation were removed [34], or violations in the distances of C*α*-C*α* (*>*4Å) and/or peptide bonds (*>*2.5Å) cutoff. The final filtered synthetic dataset used 3.5K structures that satisfied all the aforementioned criteria, and was used for pre-training the Nanobody model (“filtered SD”) before finetuning on experimental structures.

## APPENDIX D Structure prediction performance

Because training every configuration to full convergence would require prohibitive GPU resources, all Nanobody model variants are trained for the same number of optimisation steps on identical hardware. Our comparisons should therefore be interpreted as evaluating sample efficiency and performance under a fixed compute budget, rather than as the converged performance of each data regime.

Below we report detailed RMSD statistics across different nanobody variable regions (AHo definition) for each of the models we evaluated.

